# Arrayed *in vivo* barcoding for multiplexed sequence verification of plasmid DNA and demultiplexing of pooled libraries

**DOI:** 10.1101/2023.10.13.562064

**Authors:** Weiyi Li, Darach Miller, Xianan Liu, Lorenzo Tosi, Lamia Chkaiban, Han Mei, Po-Hsiang Hung, Biju Parekkadan, Gavin Sherlock, Sasha F Levy

**Author notes:** Co-first authors (Weiyi Li, Darach Miller, Xianan Liu).

## Abstract

Sequence verification of plasmid DNA is critical for many cloning and molecular biology workflows. To leverage high-throughput sequencing, several methods have been developed that add a unique DNA barcode to individual samples prior to pooling and sequencing. However, these methods require an individual plasmid extraction and/or *in vitro* barcoding reaction for each sample processed, limiting throughput and adding cost. Here, we develop an arrayed *in vivo* plasmid barcoding platform that enables pooled plasmid extraction and library preparation for Oxford Nanopore sequencing. This method has a high accuracy and recovery rate, and greatly increases throughput and reduces cost relative to other plasmid barcoding methods or Sanger sequencing. We use *in vivo* barcoding to sequence verify >45,000 plasmids and show that the method can be used to transform error-containing dispersed plasmid pools into sequence-perfect arrays or well-balanced pools. *In vivo* barcoding does not require any specialized equipment beyond a low-overhead Oxford Nanopore sequencer, enabling most labs to flexibly process hundreds to thousands of plasmids in parallel.

## Introduction

Sequence verification of plasmid DNA is a cornerstone of many cloning and molecular biology workflows. Sanger sequencing, a method developed in 1977 (1) and commercialized in 1986, remains one of the most commonly used methods. In its current form, Sanger sequencing uses a DNA primer, a DNA polymerase, and fluorescent dideoxy chain terminating nucleotides to produce DNA fragments, which are then separated by capillary electrophoresis to “read” which nucleotide caused a termination at each position. The method requires a relatively clean plasmid template (e.g. a miniprep) or amplicon and results in reads of <1 kilobase for ∼$2-5/reaction, not including the cost of DNA purification (∼$2) or a sequencing primer (∼$8). Although essential for small-scale studies, Sanger sequencing can be expensive and time-intensive for applications that require sequencing many plasmid clones or long DNA molecules: each clone requires a separate plasmid purification and each ∼500-600bp region of a long DNA molecule requires a separate sequencing primer and Sanger reaction. Because of this, alternative methods that leverage high-throughput sequencing technologies are being developed.

*In vitro* methods have been developed to multiplex high-throughput sequencing by introducing DNA barcode “indices” via PCR (2–5) or Tn5 transposase tagmentation (6, 7) (see also https://www.octant.bio/blog-posts/octopus-v3), enabling the Illumina or Oxford Nanopore Technology (ONT) sequencing platforms to sequence many barcoded samples at once. Illumina provides the highest throughput and accuracy at the lowest cost, but maximum read lengths of 125-300bp make barcoding and read assembly schemes more challenging and can miss some common plasmid features such as long repeated elements, structural variation, or plasmid multimers (8, 9). By contrast, the ONT platform generates long reads spanning entire plasmids at reasonable quality and offers better discrimination between these long DNA features (10, 11). Additionally, the ONT instrument is inexpensive enough to be purchased and run by most academic labs, and the ability to vary the user-defined runtime enables labs to scale the sequencing throughput and cost to meet variable demand. This flexibility means that a high-throughput plasmid barcoding and sequencing method built on the ONT platform could be more widely accessible. Indeed various *in vitro* protocols (3, 7) and commercial services (https://www.plasmidsaurus.com and https://www.primordiumlabs.com) have been developed to more efficiently multiplex ONT plasmid sequencing. However, all these methods require that each plasmid is purified from cells and barcoded individually, with these library preparation steps having an outsized impact on the overall cost and throughput of plasmid sequencing.

Here, we develop a **B**acterial **P**ositioning **S**ystem (**BPS**): a platform that uses bacterial conjugation, *in vivo* DNA cutting, and *in vivo* recombination to barcode and index plasmids in large bacterial arrays. This platform enables different samples to be pooled before plasmid isolation, library preparation, and ONT sequencing, greatly increasing throughput of routine plasmid sequencing. We show that BPS can sequence, with high accuracy and recovery rate, tens of thousands of plasmids in parallel at a cost between $0.12 and $1.40 per plasmid (a 5- to 70-fold cost reduction relative to existing protocols). To demonstrate new capabilities that come with this increased scale, we show that BPS can be used to transform overdispersed error-containing oligonucleotide and gene library pools into sequence-verified arrays and well-balanced pools.

## Materials and Methods

### BPS protocol for *in vivo* barcoding and pooled sequencing of plasmid arrays

For the most current version of a “wetbench” protocol, visit http://darachm.gitlab.io/bps/ and navigate to the “BPS Protocol”. For questions, suggestions, or issues, please open an Issue at http://gitlab.com/darachm/bps/-/issues.

### Bacterial strains

The donor strain used for all experiments was BUN20 [Δlac-169 rpoS(Am) robA1 creC510 hsdR514 ΔuidA(MluI):pir-116 endA(BT333) recA1 F’(lac+ pro+ ΔoriT:tet)] (12). The recipient strain was BW28705 [lacIQ rrnB3 ΔlacZ4787 hsdR514 Δ (araBAD)567 Δ (rhaBAD)568 galU95 ΔendA9:FRT ΔrecA635:FRT] (36). Both strains were kind gifts from Stephen Elledge.

Cloning and propagation of the donor plasmids was performed in BW23474 [Δlac-169 rpoS(Am) robA1 creC510 hsdR514 ΔuidA(MluI):pir-116 endA(BT333) recA1] (37). DH5α and DH10β were used for cloning and propagation of recipient plasmids. (Table S3)

### Media and chemicals

Luria-Bertani (LB) broth as a complex medium was routinely used for cloning and for growth of donor and recipient plasmids. To maintain plasmids, antibiotics were added at the following concentrations: kanamycin (Kan) (50 μg/mL), spectinomycin (Spe) (50 μg/ml), nourseothricin (clonNat) (100 μg/mL), hygromycin (Hyg) (50 μg/mL for donor plasmids and 200 μg/ml for recombinant plasmids), gentamicin (Gen) (20 μg/mL). Hygromycin is salt sensitive, so the sodium chloride for media containing hygromycin was set to 5g/L. IPTG (0.4 mM), L-arabinose (0.2% w/v) and L-rhamnose (0.2% w/v) were used to induce the P_lac_, P_araBAD_ and P_rhaBAD_ promoters, respectively. Glucose (2% w/v) was used to suppress the P_rhaBAD_ promoter.

### Construction of barcoded donor plasmids and arrayed donor clones

The backbone of donor vectors (pSL438 and pSL439, Table S4) were constructed to contain 1) KanR (kanamycin resistance), 2) oriT (origin of transfer), 3) R6K oriγ (conditional replication origin depending on the phage-derived pir1 expression), and 4) a swapping region, I-SceI-HU-HD-I-SceI, where I-SceI is the recognition site of the endonuclease SceI, and HU (5’-ttgccctctctcttcattcagggtcatgagaggcacgccattcaaggggagaagtgagatc-3’) and HD (5’-aagaacttttctatttctgggtaggcatcatcaggagcagga-3’) are the upstream and downstream homology regions for recombination. In the swapping region, a selection cassette containing HygR-SacB was cloned between HU and HD. To insert random barcodes into donor backbones (pSL438 and pSL439), an oligonucleotide library (pXL633, Table S5) that contains a NotI restriction site, a barcode region including a random 15 nucleotides, and a region of homology to both donor backbones, was ordered from IDT. pXL633, paired with pXL585, was used to PCR the barcodes with ∼1 ng of either pSL438 or pSL439 as template. The resulting PCR products were restriction digested and ligated into the corresponding donor vector via NotI and XmaI sites. Following the same cloning protocol above, the ligation products were transformed into competent BUN20 donor cells and the barcoded donor clones were selected on the LB + Kan agar plates at 37°C. Transformant clones were then randomly selected and arrayed to generate two 96-well barcoded donor collections: pSL438_BC and pSL439_BC (Table S4). To identify the barcode sequences in the arrayed donor collections, the regions containing the barcodes were amplified by colony touch PCR using pXL583 and pXL584 (Table S5) as primers. The amplicons were then purified and Sanger sequenced using pXL583. Barcodes were then extracted to compile two lists of donor barcode collections.

### Construction of barcoded recipient plasmids and arrayed recipient clones

Plasmid pSL937, which is used as the backbone to insert the random barcodes to generate the arrayed and barcoded recipient collection, was constructed from the following sources: 1) plasmid backbone/origin of replication from pBR322 (38, 39), 2) GmR (gentamicin resistance marker) from pUC18-mini-Tn7T-Gm (40), 3) homology sequences HU and HD, and two I-SceI recognition sites in a HU-I-SceI-I-SceI-HD configuration, 4) a rhamnose-inducible toxin relE (P_rhaBAD_-relE) from pSLC-217 (14), which was cloned between two SceI sites. To insert random barcodes into the recipient backbone (pSL937), an oligonucleotide library (pXL631, Table S5) that contains an XhoI restriction site, a barcode region including 20 random nucleotides, and a region of homology to pSL937, were ordered from IDT. pXL631, paired with pXL154 (Table S5), was used to generate barcodes via PCR with ∼1 ng pSL937 as template. The resulting PCR products were digested and ligated into pSL937 using MluI and XhoI restriction sites. The ligation products were then transformed into competent BW28705 cells that contain a spectinomycin-resistant helper plasmid pSL361, which was constructed by integrating a multi-cloning site and the pBAD-I-SceI (Addgene) endonuclease gene into pML104 (Table S4). Barcoded recipient clones were selected on the LB + Sp + Gm + 2% Glucose at 30°C. Transformants were then randomly selected and arrayed into 10 96-well plates.

The previously constructed two donor barcode plates were repeatedly used to acquire position and sequence identity of unknown barcodes on recipient plasmids to construct recipient positioning barcodes plates. Because sequencing chimeras may generate and mis-associate donor barcodes to recipient barcodes, each of these 96-array recipient barcode plates were mated with two donor plates to allow cross-validations and ensure accurate parsing results. The resultant recombinants containing known donor barcodes and unknown recipient barcodes were sequenced by Illumina MiSeq platform. A total of 831 unique recipient barcodes were detected and 768 (requiring a pairwise hamming distance > 5) were randomly rearrayed into 8× 96-array and 2× 384-arrayed plates, which serve as positioning barcode plates for parsing unknown DNA blocks in donor plasmids.)

### Construction of donor plasmid libraries containing oligonucleotide pools and arrayed donor clones

pSL1071 and pSL1064, which contains the NsrR-PheS or HygR-SacB cassettes, respectively, two I-SceI sites, and two homology regions for recombination (HU and HD), were used as the backbone to insert oligonucleotide pools. The 300-base oligonucleotide pools were ordered from IDT and Twist according to the following design,

GCTTATTCGTGCCGTGTTATGGCGCGCCNN…NNGCGGCCGCGGGCACAGCAATCAAAAGTA, where

GCTTATTCGTGCCGTGTTAT and GGGCACAGCAATCAAAAGTA are priming sites for the forward and reverse primers (skpp-101-F and skpp-101-R, Table S5) to amplify the oligonucleotide pool, GGCGCGCC and GCGGCCGC are recognition sites for restriction enzymes AscI and NotI, and NN…NN is the 244-base sequences randomly selected from the human genome assembly (GRCh38) by using a custom python script (Table S6). The amplification of the oligonucleotide pool was performed with ∼5 ng of template DNA and KAPA HiFi polymerase (Roche) with the annealing temperature at 53°C and extension time of 15 sec for 14-20 cycles. PCR products were purified using DNA Clean & Concentrator-5 (Zymoresearch). To clone PCR products into the donor plasmid pSL1071 or pSL1064, AscI and NotI restriction enzymes were used. The digestion reactions of PCR products and pSL1071/1064 were performed at 37°C for 4 hours. Digested products were then size selected by running a 1.2% agarose gel and recovered using Zymoclean Gel DNA Recovery Kit (Zymoresearch). The ligation reaction was performed with 25ng of digested vector and 3.8ng of inserts using T4 DNA ligase (NEB) at 16°C for 15 hours. Donor plasmids were then transformed into the donor cells by heat shock at 42°C for 60 sec and recovery at 37°C for 45 min. Resulting donor clones were randomly arrayed on PlusPlates. 384-arrayed donor clones were generated by PIXL colony picker.

### Pooled amplicon sequencing on the Illumina platform

To extract the recombinant plasmids, cells were scraped from each 96-position array selection plate and a pooled plasmid extraction was performed using Plasmid Plus Mini Kit (QIAGEN). The plasmid DNA was quantified and diluted to ∼ 1ng/μL, which is approximately 1.5×10^6^ copies of each unique barcode-barcode pair per 96-array plate. A two-step PCR was performed, as described (41) with modifications. First, 4-5 cycles of PCR with OneTaq polymerase (New England Biolabs) was performed using the forward (pBPS_fwr) and reverse (pBPS_rev) primers listed in Table S9. ∼1 ng of recombinant plasmid DNA was amplified in a single 50 μL PCR reaction with the annealing temperature at 55°C and extension time of 20 sec. To increase the multiplexing of sequencing samples, a unique pair of 1st and 2nd PCR primers (see Table S9) were used to amplify the plasmid DNA from each mated plate, which uniquely barcodes each amplification reaction and enables pooling of multiple mated plates together in one sequencing library.

Primers for the first step PCR have this general configuration and sequences are listed in Table S9:

pBPS_fwr: ACACTCTTTCCCTACACGACGCTCTTCCGATCTNNNNNNNNXXXXXXttcggttagagcggatgtg

pBPS_rev: GTGACTGGAGTTCAGACGTGTGCTCTTCCGATCTNNNNNNNNXXXXXXXXXaggtaacccatatgcatggc.

The Ns in these sequences correspond to any random nucleotide and are used in the downstream analysis to remove skew in the counts caused by PCR jackpotting. The Xs correspond to one of several multiplexing tags, which allows different plasmid pools to be distinguished when loaded on the same sequencing flow cell. The lowercase sequences correspond to the priming sites on the recombinant plasmids. The uppercase sequences correspond to the Illumina Read 1 or Read 2 sequencing primer. The PCR products were purified using NucleoSpin columns (Macherey-Nagel) and eluted into 33 μL water. A second 23-25 cycles PCR was performed with PrimeStar HS polymerase (Takara) or KAPA HiFi DNA Polymerase (Roche), with 33 μL of cleaned product from the first PCR as template and 50 μL total volume per tube. The annealing temperature is 69°C and extension time is 20sec. Primers for this reaction were the standard Illumina TruSeq dual-indexed primers (D501-D508 and D701-D712) listed in Table S9.

PCR products were cleaned using NucleoSpin columns. Amplicons from each mating plate were uniquely labeled with our customized primer indexes (first step PCR) as well as standard Illumina indices (second step PCR). This quadruple-indexed strategy increases the multiplexing capacity for sequencing. Cleaned amplicons were pooled and paired end sequenced at ∼800 reads per barcode-barcode pair on an Illumina MiSeq, HiSeq or NextSeq with 25% PhiX genomic DNA spike-in.

### Illumina sequencing data analysis for demultiplexing and sequence verification

Donor-recipient double barcode amplicon sequencing data was analyzed by customized Python scripts and Bartender. First, Illumina reads were demultiplexed using the Illumina indices. Any sequence without an exact match to two Illumina indices was discarded. Barcodes were extracted from demultiplexed sequences using the regular expressions “\D*?(.GGC|T.GC|TG.C|TGG.)\D{4,7}?AA\D{4,7}?TT\D{4,7}?(.CGG|G.GG|GC.G|GCG.)\D*” (donor barcode) and “\D*?(.ACA|G.CA|GA.A|GAC.)\D{4,7}?AA\D{4,7}?AA\D{4,7}?TT\D{4,7}?(.TCG|C.CG|CT. G|CTC.)\D*” (recipient barcode). Unique molecular identifiers (UMIs, the Ns in pBPS_fwr and pBPS_rev) were also extracted based on their expected position in the Illumina reads. Barcode reads, which contain a mix of true barcode sequences and sequences that contain errors stemming from PCR or sequencing, were next clustered into consensus sequences using Bartender (42). Each barcode cluster was next examined for replicate UMIs (indicating PCR duplicates) using Bartender, and all duplicates were removed to generate final counts of each barcode pair. The double barcodes with less than 20 reads were excluded, many of which are expected to be PCR chimeras (barcodes fused by PCR amplification). The remaining reads were used to ascertain the position of each donor barcode from each corresponding recipient barcode. Customized scripts are available at https://github.com/Li-WY/BPS-data-analysis.

### *E. coli* pre-LASSO probe design

The pre-LASSO probes (∼160-180 nt) used in this study were designed from the *E. coli* str. k-12 substr. mg1655 reference ORFeome (RefSeq: NC_000913.3) by using a custom bio-python script. The algorithm was set up to select probes that capture *E. coli* ORFs ranging from 999 bp to 2000 bp in size. The ligation and extension arms of pre-LASSO probes had similar melting temperatures, in the 65-70°C range (43).

The complete list of the pre-LASSO probes with the targeted ORFs are included in Table S8. The 417 pre-LASSO probes were obtained as pooled oligonucleotides from Twist Bioscience and used for the assembly of mature LASSO probes, as described (43). The pre-LASSO design was: 5’ CAGACGACGGCCAGTGTCGAC, Ligation Arm, AACACTTCTTGCGGCGATGGTTCCTGGCTCTTCGATC, Extension Arm, GGATCCTACGGTCATTCAGC 3’.

The assembly of the LASSO probes was performed by using a 350bp backbone: 5’TCGAGGAATTCAGAGAAGTCATCAAAGAGTTTAAAGAGTTTATGAGATTTAAGGTCAAGACAACGAGACACGAGTTCGAGATTGAGGGAGAGAAGGCCCCTCAGCGGCCTTATAACTATAACGGTCCTAAGGTAGCGAACGAACAAACCGCTAAGCTCAAGGTCACAAAAGGTCGACGAGGACCCGGATCCCTCCCCTTCTCCTGGTACGGAAGCAAAGCCTATGTTAAACACTGACTATCTGAAGCTCTCCTTCCCTGAAGGCTTGAGAGATTCATGAACTTCGAGGAAGGACGGAGAGTTTATTTATAAGGAACCAACTTCCCCTCCGATGGCCCTGTCATGAATTCT 3’

### Capture and cloning of *E. coli* ORFs by LASSO probes

Capture of *E.coli* ORFs was performed as described (44). Briefly, the 417 LASSO probe library was hybridized to *E. coli* genomic DNA in 15 μL of 1× Ampligase DNA Ligase Buffer (Epicentre) containing 250 ng of unshared *E. coli* K12 total genomic DNA and 5 ng of the LASSO probe pool. In the hybridization reaction, the concentration of *E. coli* chromosomes was approximately 10 pM. The reaction (15 μL) containing the LASSO probe pool and the *E. coli* genomic DNA was denatured for 5 min at 95 °C in a PCR thermocycler (Eppendorf Mastercycler), then incubated at 65 °C for 60 min. After hybridization, 5 μL of freshly prepared gap filling mix (Table S7) was added into the hybridization solution while maintaining the reaction at 65°C in the thermal-cycler. Gap filling and ligation was performed for 30 min at 65°C. After capture, the DNA samples were denatured for 3 min at 95°C, and the temperature was reduced to 37°C. Next, 4 μL of linear DNA digestion solution was added. Digestion was performed for 1 h at 37°C, followed by 20 min at 80°C.

A post-capture PCR was performed using AttB1CaptF and AttB1CaptR (Table S5) as described (43). The post capture PCR product was purified by using AMPure XP Beads (Beckman Coulter) and mixed with the Gateway ‘donor vectors’ (pDONR221, (45)) and the BP Clonase enzyme mix (Invitrogen). The BP reaction was purified and used for electroporation in NEB 10-beta Electro-competent *E. coli* (c3020K) to generate cloned libraries.

### Integration of captured ORF pools into BPS donor plasmids

Plasmid pSL1064, which contains the HygR-SacB cassette, two I-SceI sites, and two homology regions for recombination (HU and HD), was used as the backbone to insert pooled *E. coli* ORFs. The *E. coli* ORFs were integrated into the plasmid pDONR221 (45). oSL1581 and oSL1582 (Table S5) were used to introduce AscI and NotI recognition sites to these ORFs for cloning into pSL1064 by PCR. The amplification of the pooled ORFs was performed with 10 ng of template ORFs and KAPA HiFi polymerase (Roche) for 20-cycle PCR with the annealing temperature at 57°C and extension time of 3.5 min. Amplified ORFs were purified using DNA Clean & Concentrator-5 (Zymoresearch). To clone amplified products into the donor plasmid pSL1064, AscI and NotI restriction enzyme recognition sites were used. The digestion reaction of amplified products and pSL1064 were performed at 37°C for 4 hours. Digested products, ranging from 1-2kb, were size selected by cutting bands from a 1.2% Agarose gel and isolating DNA using Zymoclean Gel DNA Recovery Kit (Zymoresearch). The ligation reaction was performed with 100 ng of digested vector backbone and 70.8 ng of inserts using T4 DNA ligase (NEB) at 16°C for 15 hours.

### Arrayed mating for barcoding DNA with BPS

Each donor plate was mated to each barcoded recipient plate in an arrayed format on agar plates. The donor arrays were grown on LB + Kan + Hyg/clonNat plates overnight at 37°C; the recipient arrays were grown on LB + Sp + Gm + 2% Glucose overnight at 30°C. The agar media for arrayed mating was LB+Ara + IPTG, pre-warmed in 37°C for 1 hour. Both donor and recipient clones were transferred onto the mating plates using SINGER ROTOR HDA robot with 96-or 384-position pin pads, and grown for 3-6 hours at 30°C. The mated cells were then transferred onto the selection plates containing LB + Ara + Rha + Gm + Hyg/clonNat. Recombinant clones were then selected at 37°C overnight.

### Pooled sequencing of whole plasmids on the Oxford Nanopore platform

Bacterial clones on selection plates were scraped and pooled to extract recombinant plasmids containing the recipient barcodes and DNA blocks (donor barcodes, oligonucleotides, and *E. coli* ORFs) using Plasmid Plus Mini Kit (QIAGEN). The pooling capacity is determined by the number of unique positioning barcodes. In this study, 768 positional barcodes were routinely used and up to 768 clones (2× 384-array plates) containing unique positional barcodes can be pooled.

Two fragmentation approaches were used to generate linearized plasmids for nanopore library constructions. One is to use the restriction enzyme PmlI (NEB) to cut circular plasmids by incubation at 37°C for 2 hours. Linearized plasmids were size selected by running a 1.2% Agarose gel and recovered using Zymoclean Gel DNA Recovery Kit (Zymoresearch). The second approach is to tagment circular plasmids using the transposome complex from Rapid Barcoding Kit (SQKzRBK114.96).

The Native Barcoding Kit (SQK-NBD112.96 and SQK-NBD114.96, Nanoporetech) and Rapid Barcoding Kit 96 V14 (SQK-RBK114.96, Nanoporetech) were used to construct sequencing libraries for the Oxford Nanopore platform. Reads were generated from MinION and PromethION flow cells (FLO-MIN112/FLO-MIN114/FLO-PRO114M, Nanoporetech).

### BPS analysis pipeline

A bioinformatics pipeline was devised to achieve appropriate, performant, and scalable analysis of BPS experiments, and this pipeline uses a flexible configuration interface to enable diverse applications. The aim of the pipeline is to (1) gather long-read sequencing data from multiple runs, (2) extract a small barcode from each read, (3) use this small barcode to separate reads from each colony, (4) perform one of two assembly strategies, and (5) assess the “purity” and “correctness” of the assembly at that position.

Basecalled FASTQ files are filtered for size and a known contaminant file is used to optionally filter out reads using alignment (minimap2 (46) and samtools (47)). Within each sequencing library pool, demultiplexing barcodes that are introduced during ONT library preparation are used to assign sample membership to each read. Alignment to unique “signature” sequences are used to separate out different plasmids within each sample (minimap2 and awk). Alignment to known “trimming” sequences is used to remove most of the backbone (minimap2 and python), and fuzzy regular expressions are used to extract exactly the barcode from a known sequence context (itermae, https://gitlab.com/darachm/itermae/). Barcodes are clustered and assigned to an optionally provided list of known barcodes (starcode (48) and python), then each combination of sample and position barcodes is used to separate raw FASTQ records into separate files (awk). These may be assembled using one of two options (A or B). (A) To assemble long (>1kb) target sequences, read-length distributions per well are analyzed with a gaussian-mixture model to separate different species of plasmid in each well (python), then each cluster of reads is used for *de novo* assembly (trycycler (15) and flye (16)) and polishing of the assembly (medaka). (B) To assemble of short (<1kb) targets, raw reads are trimmed using alignment to known trimming sequences (minimap2 and python) before multiple-sequence alignment (kalign3 (49)), merging to a draft consensus (python), and polishing of the consensus (racon (50) and medaka). The purity of either assembly is assessed by aligning each raw input sequence to the resulting assembly (either with minimap2 (46) or with pairwise alignment using BioPython (51)). The resulting assemblies are subject to another round of post-assembly processing using options of raw output, a “rotation” to begin all of the circular assemblies in a similar location (minimap2 (46) and python), trimming of known sequences (minimap2 (46) and python), and/or exact extraction using fuzzy regular expressions (itermae). Final processed assemblies are optionally compared to a known target sequence file to assess “correctness” (minimap2 (46) or bwa (52)). Each successfully considered position is analyzed (using R (53)) to output a per-position per-sample call of purity and correctness (in matching a particular reference in the provided set). The entire pipeline is written in Nextflow (54), uses Singularity (55) to execute Docker containers, and makes extensive use of GNU utilities (including GNU parallel (56)).

The pipeline is available on GitLab under the BSD 3-clause license, documentation is available at darachm.gitlab.io/bps, and we will support users via Issues at gitlab.com/darachm/bps-dev/-/issues.

### Illumina sequencing to evaluate the uniformity of oligonucleotides before and after amplification

To approximate the abundance of the original single stranded oligonucleotides generated from service providers, second strands were synthesized using a primer annealing and extension approach (57–61). The second strand synthesis reaction was performed with ∼50 ng (IDT) or ∼8 ng (Twist) of template DNA and KAPA HiFi polymerase (Roche) with the annealing temperature at 53°C and extension time of 30 sec for 2 (IDT) or 10 (Twist) cycles. For second strand synthesis only one primer (skpp-101-R) was used.

The resultant dsDNAs, together with amplicons of oligonucleotides generated from 14 (IDT) or 20 (Twist) cycles of PCR, were ligated with xGen™ UDI-UMI Adapters (IDT) which contain full length sequencing primers, i7/5 indices, and Unique Molecular Identifiers (UMIs). PE150 reads were generated by Illumina iSeq platform. The absolute copy number of oligonucleotides/amplicons were estimated by counting the number of unique UMIs associated with each type of oligonucleotide/amplicon.

## Results

### Overview of the *in vivo* barcoding platform

MAGIC cloning (12), developed as an *in vivo* alternative to *in vitro* GATEWAY cloning (13), enables rapid subcloning of a DNA block from one plasmid to another using bacterial conjugation and *in vivo* recombination. A DNA block in a donor plasmid is conjugated to a recipient cell containing a recipient plasmid. A genetic program in the recipient cell recombines the DNA block from the donor plasmid to the recipient plasmid using the endonuclease *SceI* to cut both plasmids and the recombinase *lambda Red* to stitch (recombine) the donor DNA block into the recipient plasmid backbone. We extended this platform for use in multiplexed plasmid sequencing by constructing arrays of cells with a barcode that is unique to a position on an array (positional barcodes) in either the donor plasmids or the recipient plasmids (Figure 1A). In addition, we made two major modifications to improve the reliability and portability of the platform. First, the *PheS* counter-selection marker is replaced by *relE*, with flanking terminators to ensure deliberate control of expression (14). Second, the homing endonuclease *I-SceI* is placed in a helper plasmid (pSL361) with a temperature-sensitive replication origin (pSC101 -ori^ts^), providing a convenient way to cure (remove) the helper plasmid prior to isolation of recombinant plasmid DNA.

**Figure 1.**
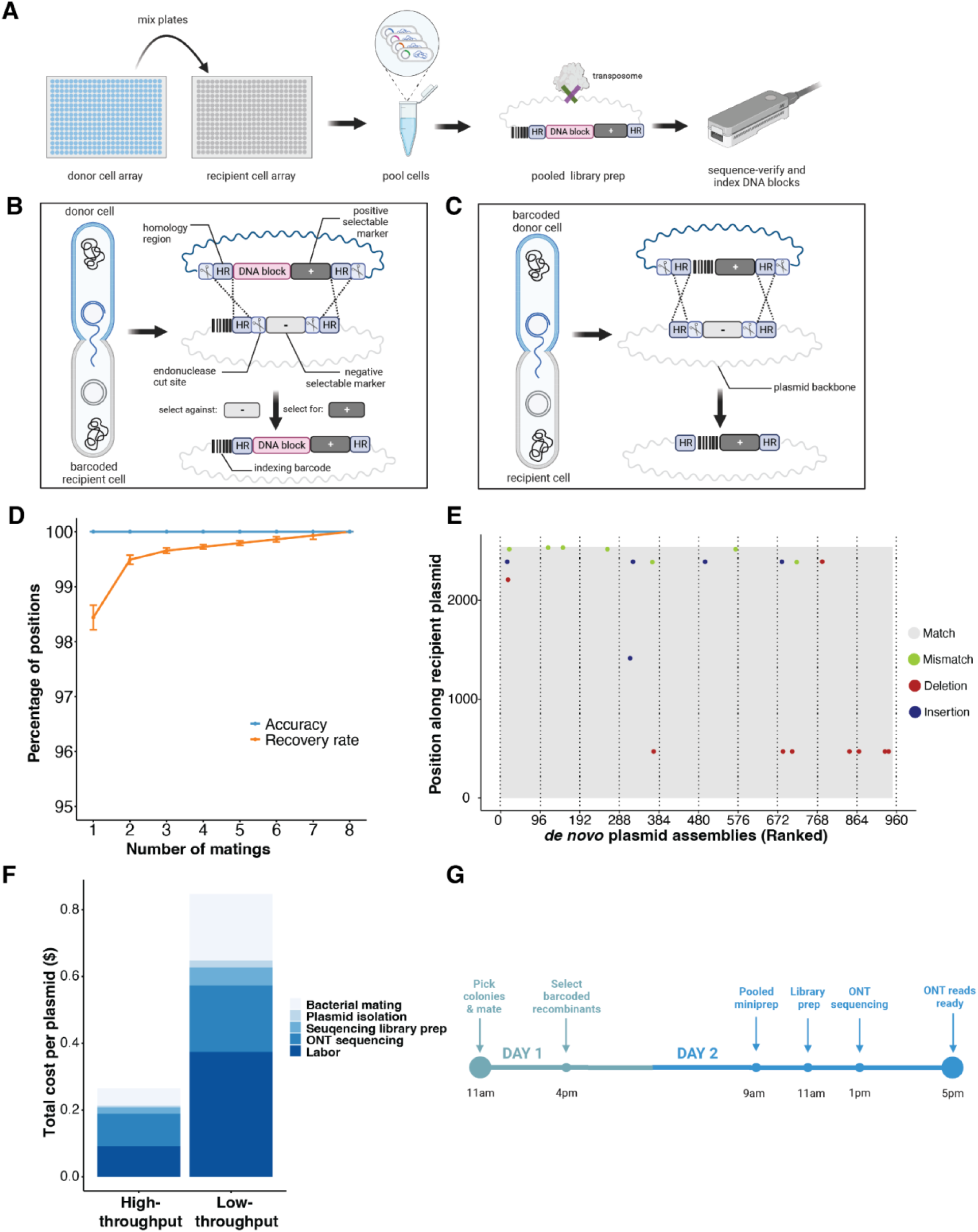
Workflow and performance of BPS, an *in vivo* barcoding platform. (A) Donor cell arrays (blue) containing donor plasmids are conjugated to recipient cell arrays (gray) containing recipient plasmids. A DNA cassette from the donor plasmid is recombined into the recipient plasmid backbone to join an indexing DNA barcode with a sequence of interest. Cells from one or more plates can subsequently be pooled and prepared for sequencing. (B) In one design, a cassette with a DNA block on a donor plasmid is recombined into a recipient plasmid backbone containing a positional barcode. (C) In another design, where a whole plasmid backbone needs to be sequenced, a donor positional barcode is recombined into a recipient plasmid. In both (B) and (C), a scissor icon is an SceI cut site, HR is a region of sequence identity that mediates homologous recombination, dashed lines are homologous recombination events, and + or - icons are selection or counter-selection markers. (D) The recovery rate (percent of positions that were detected, orange) and accuracy (percent of detections that were sequence correct, blue) for the design in (B). Number of matings (x-axis) indicates the number of times the same DNA block was mated to a barcode and sequenced. Error bars indicate standard errors calculated by bootstrapping. (E) In the design illustrated in (B), donor barcodes were mated to recipient plasmids and plasmid sequences at each position were assembled *de novo* by sequencing pools of clones. Variation from the *a priori* reference expectation, including insertions, deletions, and substitutions are shown by colored dots. Successful *de novo* assemblies are ranked by decreasing ONT read coverage. (F) The cost of plasmid sequencing when BPS is performed at high- and low-throughput. Low-throughput assumes barcoding is performed in 96-well plates and 4,608 plasmids are sequenced at 100-400× coverage per flow cell. High-throughput assumes barcoding is performed on 384-position agar arrays and 9,216 plasmids are sequenced at 100-200× coverage per flow cell. Detailed cost assumptions are listed in Tables S1 and S2. (G) Experimental timeline for sequence verification by BPS. We assume a Minion flow cell contains 250 active pores generating data at 100 bases/second, enabling ∼500 (7kb) plasmids to be sequenced at 100x depth in 4 hours.

In one design (Figure 1B), DNA constructs are sequenced by integrating them into donor plasmids that are subsequently conjugated to arrays of cells containing recipient plasmids with positional barcodes. In another design (Figure 1C), the whole backbone of recipient plasmids are sequenced by conjugating positional barcodes from arrays of donor cells and recombining the barcodes into the recipient plasmids. The resultant recombinant plasmids from either workflow, including positional barcodes and to-be-sequence-verified plasmid DNA, can be pooled and processed for ONT sequencing with a single DNA miniprep and a single ONT library prep. To process these data, we developed a flexibly-configured Nextflow-based pipeline to enable read partitioning and scalable *de novo* assembly of tens of thousands of plasmids.

### Accuracy and Recovery Rate

To test the accuracy and recovery rate of the *in vivo* barcoding and sequencing platform, we generated 96-position arrays of 192 clones of barcodes in donor cells/plasmids and 768 clones of barcodes in recipient cells/plasmids, verified the barcode sequence at each position using a combination of Sanger and Illumina sequencing (Materials and Methods), and used these arrays as a test set. We next mated each donor array to each recipient array (768 matings per donor array), selected for colonies that contain barcode-barcode recombinant plasmids, and performed two minipreps (one for each donor array mating). For each plasmid pool (768 clones), we tagmented plasmids using the ONT Rapid Barcoding Kit 96 (V14). Each tagmentation reaction introduces one of 96 unique sample indices to the plasmid pool enabling up to 73,728 (96 × 768) plasmids to be processed in parallel with our 768-recipient barcode array. We sequenced plasmid pools on an ONT Minion (R10.4.1) at average sequencing depth of 130 reads per position. A custom BPS bioinformatics pipeline (Materials and Methods) was developed to partition ONT sequencing data by both the positional barcodes (introduced *in vivo* via conjugation) and the sample indices (introduced *in vitro* during ONT library preparation), and to assemble (using Trycycler (15) and Flye (16)) and polish (using medaka https://github.com/nanoporetech/medaka) the consensus sequence for each position on each 96-position array. Consensus ONT sequences of barcode-barcode pairs in all cases matched the sequence determined by Illumina and Sanger sequencing (Materials and Methods). Using these ONT data, we calculated the recovery rate (percent of known positions that were detected) and accuracy (percent of detections that were correct) of donor barcode sequences (Figures 1D and S1). We found perfect accuracy and a high recovery rate (>98%) that improves when the same donor barcode is assayed multiple times (Figure 1D). Missing positions were, in all cases, due to a lack of sufficient sequencing coverage (Figure S2). ONT reads also enabled *de novo* assembly of the whole plasmids to detect various types of sequence variation from the recipient backbone (Figure 1E).

### Low-cost whole-plasmid sequencing with a fast turnaround time

The number of plasmids that can be processed in parallel by *in vivo* barcoding is determined by the number of unique positional barcodes across arrays. Here, we constructed arrays of 768 positional barcodes to enable DNA isolation and library construction of pools of 768 plasmids in parallel. At this scale of multiplexing, using off-the-shelf robotics for colony picking and arrayed bacterial conjugation/mating, we estimate that thousands of plasmids can be sequenced in parallel for between $0.12 (100× sequencing depth) and $0.53 (1,000× sequencing depth), including all consumables and labor costs (high throughput in Figure 1F and Table S1). At lower scales of multiplexing, we estimate that manual colony picking and mating can be used to sequence hundreds of plasmids in parallel for between ∼$1.00 (100× sequencing depth) and $1.40 (1,000× sequencing depth), including all consumables and labor costs (low throughput in Figure 1F and Table S2). Because most steps of the procedure are performed with cell or plasmid pools, both low-throughput and high-throughput protocols can be performed with ONT reads ready as soon as the next day (Figure 1G).

### Demultiplexing and sequence verification of oligonucleotide pools

Generation of oligonucleotide pools using arrayed synthesis technologies can be several orders of magnitude less expensive than one-at-a-time column-based DNA synthesis ($0.0005-0.035/base vs. $0.07-0.50/base) (17, 18). Parallelized methods that capture long blocks of natural DNA can offer similar cost savings relative to *de novo* gene synthesis. Yet, many testing modalities require arrays of DNA designs (e.g., mass spectrometry, microscopy, and enzymatic assays) and are unable to take advantage of these sources of low-cost DNA. We next explored whether the higher throughput of the BPS plasmid sequencing platform could be used to demultiplex such DNA pools.

To test the capability of BPS to demultiplex oligonucleotide pools, we designed a library of 1,100 oligonucleotides, each containing a 244-base sequence chosen randomly from the human reference genome GRCh38 (Figure 2A) (19). Differences in the efficiency of synthesis across different nucleotide sequences may result in oligonucleotide pools that are more or less dispersed in frequency (defined here as pool dispersion). Higher pool dispersion would require picking and sequencing more clones per design (higher sampling depths) to recover the same number of designs across an array (20). To minimize potential pool dispersion in our first test, we subset the 1,100 oligonucleotide designs into one 100 randomly-chosen oligonucleotide pools that were scored as “low complexity” by the IDT online oligonucleotide analysis tool (https://www.idtdna.com/site/order/plate/gblocks). This pool was synthesized by IDT as an oPool, and inserted into donor plasmids/cells by PCR, digestion, and ligation. Randomly arrayed clones were generated from this pool at a ∼20× sampling depth (the average number of clones picked per DNA block in the pool) for *in vivo* barcoding and sequencing, as described above, to generate a consensus sequence at each position on each plate. Given the relatively high error rate of ONT sequencing, we needed to distinguish positions that contain only sequencing errors from those that contain a mixture of two or more plasmids. For each position, we determined a purity score (the fraction of ONT reads that are >90% end-to-end identical to the consensus sequence at that position) and assayed, across a range of purity scores, which clones were indeed pure by examining Sanger sequencing traces (Figure S3). We found that purity scores >0.8 were reliably pure clones (15/15) but that those purity scores < 0.8 were frequently mixed clones (7/8). Based on these results, we set a conservative 0.9 purity score threshold for calling a clone “pure” and flagged putatively “impure” positions. Illumina sequencing was performed on a subset of samples to validate the correctness of consensus sequences (Materials and Methods), and for unflagged positions the consensus Illumina and ONT sequences agreed 99.7% of the time (with most differences presumably stemming from errors in PCR during the Illumina library prep). After discarding flagged positions, we recovered a sequence-perfect clone for 83% of the oligonucleotide designs in this pool.

**Figure 2.**
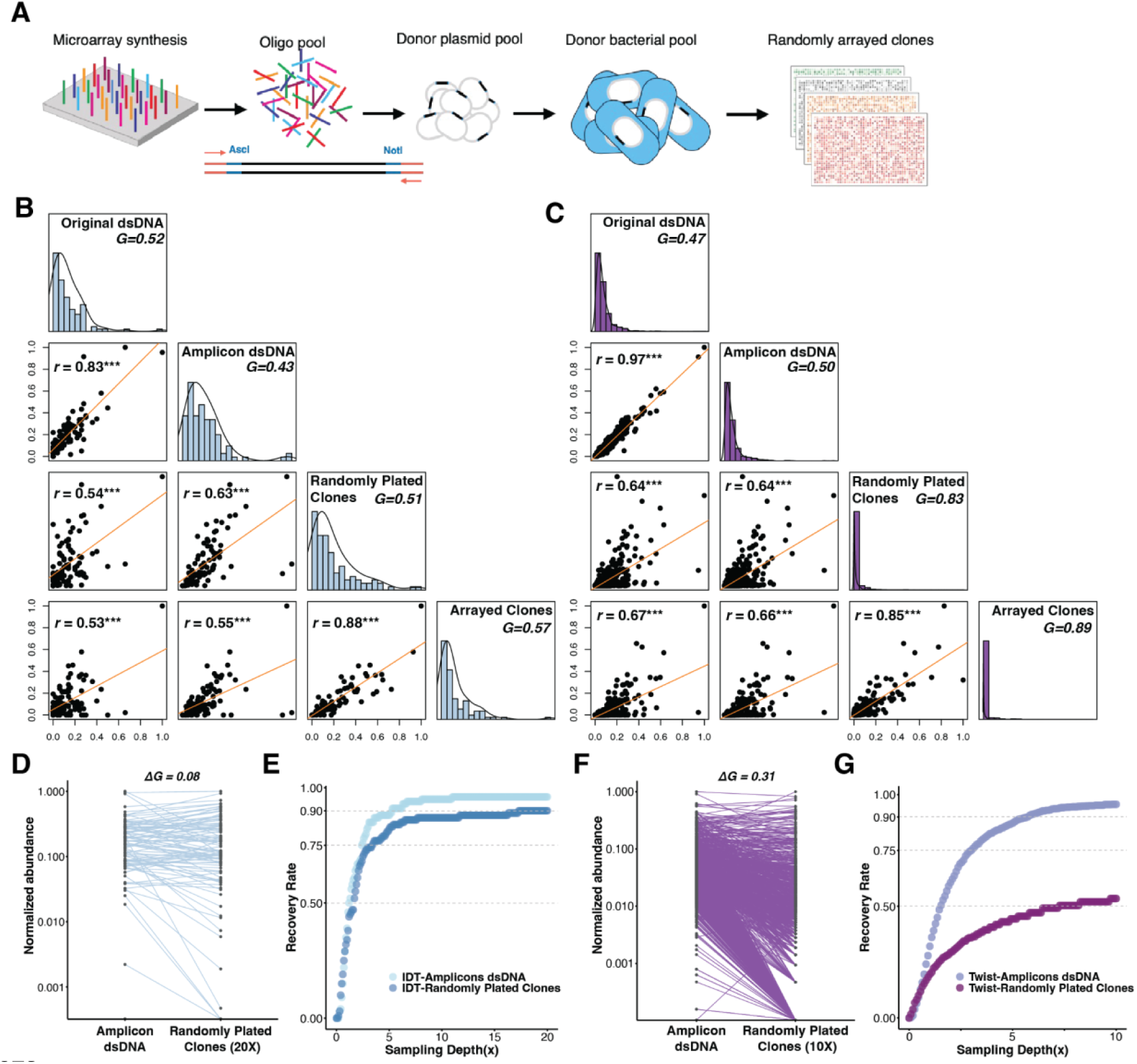
Sequence verification and demultiplexing of oligonucleotide pools. (A) Oligonucleotide pools from vendors were cloned into donor plasmids and donor cells, and randomly arrayed. For experiments here, oligonucleotides are designed with fixed priming sites (red arrows) and restriction endonuclease sites (blue, AscI and NotI) to facilitate cloning. (B) and (C) The distribution of oligonucleotide abundances from the IDT (B) and Twist (C) pools, which contain 100 and 1,100 oligonucleotide designs respectively, assessed at multiple stages during the *in vivo* barcoding workflow: 1) original oligonucleotide pools after the second strand synthesis (original dsDNA), 2) following 14-20 rounds of PCR amplification (Amplicon dsDNA), 3) random clones after inserting the oligonucleotides into plasmid backbones and cells and randomly plating for clones (Randomly Plated Clones), 4) and parsed clones after *in vivo* barcoding and sequencing (Arrayed Clones). Pairwise comparisons of the abundance of each oligonucleotide design at different stages are displayed in scatter plots with Pearson correlation coefficients (*r*). The level of corresponding significance is indicated by asterisks (\*\*\**P <* 10^-7^). Histograms represent frequency distributions of oligonucleotide designs with different abundance levels. Gini coefficients, *G*, are used to quantify the degree of pool dispersion. A uniform distribution has a Gini coefficient of 0. (D) and (F) The change of normalized abundance of each oligonucleotide design from amplicons to clones for the IDT and Twist pools, respectively. *ΔG* denotes the increase in Gini coefficients from the Amplified dsDNA to Randomly Plated Clones pools. (E) and (G) Recovery curves for all oligonucleotide designs at different sampling depths for the IDT and Twist pools, respectively. The expected recovery rate at different *in silico* sampling depths is calculated by assuming the probability of sampling one oligonucleotide design equals its empirical frequency observed in the corresponding pool.

We next scaled-up the method by demultiplexing, at ∼20× sampling depth, a pool containing all 1,100 oligonucleotide designs, synthesized by Twist. This full set includes 100 oligonucleotides that were scored as “intermediate complexity” by the IDT online oligonucleotide analysis tool, indicating that it may be more dispersed than the 100 oligonucleotide pools synthesized by IDT. We recovered sequence-perfect clones for 51.5% (567/1100) of the oligonucleotide designs in the full pool, and 52.0% (52/100) of the oligonucleotide designs in the subset with the same designs as the subpool synthesized by IDT.

### Examination of factors impacting the recovery rate of demultiplexing

Given the high recovery rate of BPS using pre-arrayed colonies as inputs (see Accuracy and Recovery Rate experiments above), we sought to understand what factors impact the recovery rate of commercial oligonucleotide pools. The recovery rate could be influenced by sequence errors introduced by 1) synthesis and/or assembly of linear DNA, 2) PCR amplification, or 3) introduction into plasmids and cells. The vendor-reported oligonucleotide synthesis error rate of ∼4 × 10^−4^/nt predicts that ∼ 11.3% (1 − 0.9996^300^) of oligonucleotides in the pool contain erroneous sequences. Additional sequence errors introduced during amplification and cloning of the BPS protocol appeared to have little impact on the recovery rate: we observed a base substitution error rate that is roughly the same as the rate reported by the vendors (4.75 × 10^−4^/nt and 3.81 × 10^−4^/nt for IDT and Twist pools respectively, Figure S4).

We next explored the impact of variation in sequence abundance on the recovery rate by examining the level of pool dispersion at different stages of the protocol for both the IDT (Figure 2B) and Twist (Figure 2C) pools: following second strand synthesis (Original dsDNA), PCR amplification (Amplicon dsDNA), introduction into plasmids and plating of cell colonies (Randomly Plated Clones), and construction of colony arrays and barcoding by BPS (Arrayed Clones). The lowest abundance correlations were observed between the Amplicon dsDNA and Randomly Plated Clones steps. Using Gini coefficients as a measure of pool dispersion, we found that this step also resulted in the greatest increase in pool dispersion, resulting from large changes in abundance of particular oligonucleotide designs (Figures 2D and 2F). To determine whether the pool dispersion at this cloning step was due to inadequate clone sampling, we estimated the expected recovery rate based on the frequency distribution we observed in the Amplified dsDNA pools (Figures 2E and 2G). We found that, at a 7× sampling depth, >93% of sequence perfect designs are expected to be present at the Randomly Plated Clones step, while we could only recover 86% and 49% for IDT and Twist pools respectively, by resampling the observed Randomly Plated Clones (Figures 2E and 2G). These data suggest that the cloning step, which includes introduction of oligonucleotide designs into plasmids and cells, has the greatest impact on frequency dispersion and therefore demultiplexing performance.

### Demultiplexing a captured open reading frame (ORF) library

To demonstrate the capability of BPS to demultiplex pools of longer DNA (1-2kb) with variable sizes, we parsed a library of ORFs captured using long-adapter single-strand oligonucleotide (LASSO) probes (21). This DNA capture technique uses pools of long inversion probes to selectively hybridize and amplify multiple target regions from genomic DNA. While LASSO probes have been demonstrated to capture >3,000 *E. coli* ORFs in parallel (21), off-target sequences are also captured, desired on-target sequences may contain mutations from PCR, and, as expected for any pooled technique, the relative abundance of each type of sequence can vary significantly. These features can limit the cost-efficiency and data quality of downstream pooled assays. To determine if BPS could be used with LASSO probes to generate ORF arrays or well-balanced sequence-perfect pools, we captured 417 *E. coli* ORFs (1-2 kb) by LASSO probes, cloned them as a pool into the BPS donor plasmid backbone, and picked a total of 10,752 clones (∼25× sampling depth) for *in vivo* barcoding and sequence verification (Figure 3A). Of the 8,562 pure clones isolated by BPS, we found that 30.7% were sequence perfect, 61.2% had at least one mismatch, and 7.5% were off-target DNA. The high number of sequences that contain at least one mismatch is likely a result of the many cycles of PCR required for the process: 25 cycles for post ORF capture amplification and 20 cycles for cloning into the BPS donor plasmids. Sequence perfect clones contained 52.3% (218/417) of targeted ORFs (Figure 3B). The distribution of DNA lengths of these error-free ORFs suggests that our protocol can capture ORFs in all size ranges in the original pool (Figure 3C).

**Figure 3.**
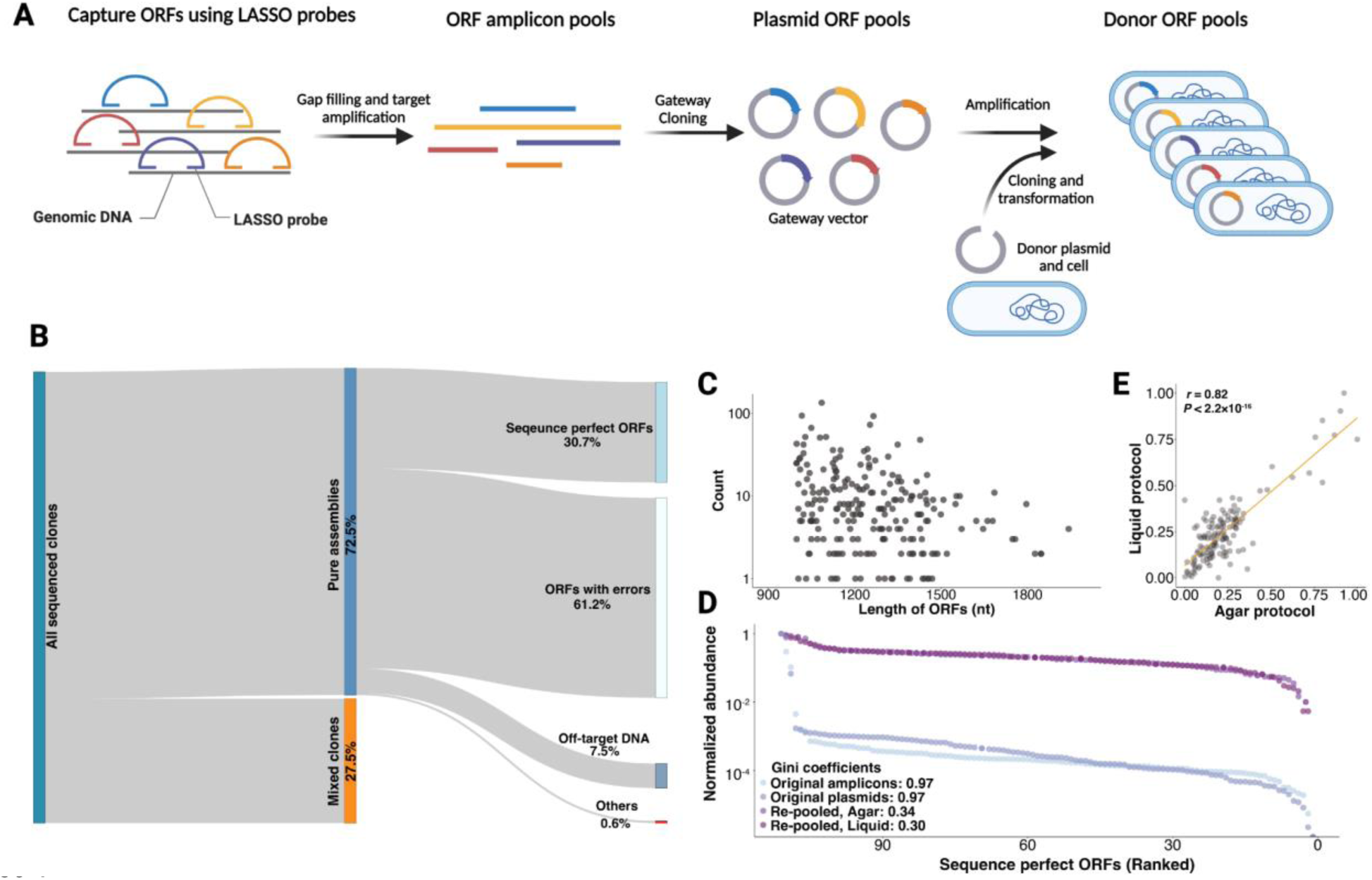
Sequence validation and construction of a balanced ORFs library. (A) Pooled capture of *E. coli* ORFs and cloning into the BPS donor cells. ORF libraries were amplified from *E. coli* genomic DNA using Long-Adaptor Single-Stranded Oligonucleotide (LASSO) probes and inserted into a Gateway vector. Then, ORFs were amplified by PCR and cloned into BPS donor plasmids and cells. (B) A Sankey diagram showing the sequencing results of the ORF library following BPS. Mixed clones (orange, purity score < 0.9, see Methods) indicate that the same position on an array is likely to contain more than one ORF. Pure clones (blue, purity score ≥ 0.9) are further classified into sequence perfect ORFs, ORFs with errors (point mutations or indels), off-target DNA (cannot be mapped to the target ORFs but can be mapped to the *E. coli* genome), and others (cannot be mapped to any region of the *E. coli* genome). (C) The length and abundance of sequence-perfect ORFs identified from pure donor clones. (D) The distribution of normalized ORF abundance in the original amplicon pools after PCRing from genomic DNA (original amplicons), after cloning into Gateway vector plasmids (original plasmids), and after re-pooling following growth of sequence perfect clones in liquid or on agar. ORFs were ranked according to their abundance from lowest to highest. Gini coefficients were calculated to quantify the degree of uniformity of ORF abundances in a pool. A lower value indicates a higher degree of uniformity. (E) The correlation of the relative abundance of each ORF from balanced libraries constructed using agar and liquid protocols.

When DNA pools contain sequence errors or are overdispersed, the power of sequencing-based pooled functional assays (e.g. massively parallel reporter assays (22)) can suffer because: 1) constructs with errors may not provide useful data; 2) more reads are needed to assay low abundance constructs; and 3) some sequence errors (such as errors result in premature stop codons) may contaminate selection schemes. To demonstrate the capability of our platform to address these issues by constructing error-free well-balanced pools, we arrayed 111 clones carrying sequence perfect ORFs, grew replicates of these arrays both in liquid multi-well plates and on agar pads, and pooled each replicate separately. For each replicate, we extracted plasmids and ONT sequenced the library (without an additional amplification). We found that demultiplexing with BPS and re-pooling drastically improved the uniformity of the pool (Figure 3D). ORFs that were highly overrepresented following LASSO capture were not overrepresented in the balanced libraries. The relative abundance of each ORF in a balanced pool was highly correlated between replicate pools using the same outgrowth procedure [Pearson’s *r* = 0.73, *P* <2.2e-16 in liquid vs. liquid, Pearson’s *r* = 0.82, *P*<2.2e-16 in agar vs. agar] or different outgrowth procedures [Pearson’s *r* = 0.82, *P* < 2.2e-16 in liquid vs. agar] (Figure 3E), suggesting that abundance differences are due to reproducible growth rate differences between clones.

## Discussion

We have developed a Bacterial Positioning System (BPS): an *in vivo* plasmid barcoding platform for high-throughput sequence validation of plasmid DNA. In contrast to *in vitro* barcoding methods that require each sample to be prepared independently (plasmid extraction and barcoding with enzymes), BPS barcodes plasmid DNA *in vivo,* enabling arrays of cells to be pooled prior to sample processing. This dramatically reduces the cost and hands-on time required, and using BPS with low-overhead ONT sequencing enables most labs to flexibly process hundreds to thousands of plasmids.

One application of high-throughput plasmid sequencing is demultiplexing of DNA pools that are the products of next-generation DNA synthesis (23), pooled DNA assembly (24–26), or pooled DNA capture (21, 27) methods. While construction of these pooled libraries is inexpensive relative to constructing each design independently, demultiplexed plasmids are required for many testing modalities (e.g., mass spectrometry, microscopy, and enzymatic assays) and for downstream DNA engineering. We have previously developed an arrayed *in vivo* barcoding platform in *S. cerevisiae* and used it to demultiplex and sequence verify pools of oligonucleotides encoding gRNA variable regions (20). However, that platform requires that both the barcode and the DNA-to-be-sequenced are placed at a similar location in the yeast genome. This limitation, in combination with the slower growth rate relative to *E. coli*, makes the yeast platform too cumbersome for most applications. Another demultiplexing solution is dial-out PCR (28–31), which uses pre-designed unique tags to prime specific sequences from a pool. Although this *in vitro* approach is expected to recover designs at low relative abundance, it is expensive and time consuming to scale up: isolation of each design requires a PCR with a unique set of primers. Scalable low-cost plasmid sequencing with technologies such as BPS offers an alternative “shotgun” approach (32–35) to demultiplexing: instead of isolating each design by bespoke methods, clones are oversampled from a pool and sequenced, with the aim of recovering a large fraction of designs.

However, shotgun demultiplexing approaches have several constraints that limit their utility. Similar to the high sequencing depths required for shotgun sequencing, the number of clones sampled (sampling depth) must be several fold the size of a library to have a good chance of recovering most designs, even when the frequency dispersion is low. Frequency dispersion was not low in some of our experiments, meaning that, even at high cloning depths, we would be unable to recover many designs (Figure 2G). Frequency dispersion can be introduced at several steps before and during the BPS protocol: DNA library construction (e.g., pooled DNA synthesis), PCR amplification, integration into a plasmid backbone, transformation into host cells, and cell outgrowth. In our experiments, we find that the measures of oligonucleotide design abundances have the lowest correlation before and after the cloning step. Several factors such as the size, initial concentration, complexity of a DNA library (Figure S5), differences in the efficiency of plasmid integration between designs, undersampling of transformants, and jackpotting of some designs following transformation may all contribute to dispersion. Despite these potential sources of variation, several groups have achieved pooled design libraries with relatively low levels of dispersion and high recovery rates (5, 24). More research is needed to determine which differences between protocols contribute to this cloning variance and how it can be minimized. Nevertheless, shotgun demultiplexing is a strategy made viable by inexpensive plasmid screening as described here, and even overdispersed pools could be made useful when an investigator only needs to sparsely sample the design space (4) (e.g. randomly sampling 10^3^ designs from a pool of 10^6^). In addition, demultiplexing and subsequent pooling of sequence-verified clones could be used to transform error-prone overdispersed pools into error-free low-dispersion pools that can be more cost-effectively assayed by sequencing readouts.

BPS provides two methods for sequencing of plasmid DNA: 1) a DNA block of interest is transferred to become adjacent to a positional barcode (Figure 1B), or 2) a positional barcode is transferred into a plasmid of interest (Figure 1C). Both methods use ONT long-read sequencing, meaning that large DNA blocks and/or plasmids with repetitive features (tandem repeats and long interspersed repeats) can be more accurately characterized than by short read methods. With the first method, DNA blocks will generally be inserted into donor plasmids using standardized approaches (e.g., the same restriction sites or homology regions for Gibson assembly), making this method attractive for routine sequence validation of DNA parts (e.g., oligonucleotides, genes, and variants) that have been synthesized, captured, or assembled. With the second method, the entire plasmid is sequenced, enabling not only verification of a DNA part of interest, but also detection of undesired changes to the plasmid backbone, such as point mutations, insertions, deletions, duplications, and rearrangements. However, plasmids sequenced by this method must have a BPS “landing pad” that accepts a barcode cassette, meaning that some backbone engineering is required prior to use with new plasmids. As an alternative, new BPS barcode cassettes could be developed to integrate at sequences that are common to most plasmids in lab use (e.g., natural landing pads). With this advance, BPS could be used on most plasmids without any modifications, dramatically reducing the cost and hands-on time for sequencing of most plasmid DNA.

## Data Availability

All sequencing data are deposited in NCBI with the BioProject Number PRJNA1026144. BPS protocol and analysis pipeline are available at darachm.gitlab.io/bps and https://github.com/Li-WY/BPS-data-analysis, and we will support users via Issues at gitlab.com/darachm/bps/-/issues.

## Supporting information

Large Supplementary Tables (S1, S2, S6, S8, and S9)

## Acknowledgements

We are grateful to Stephen Elledge for providing MAGIC strains and plasmids, to Kevin Roy, Lars Steinmetz and the Stanford Genome Technology Center for providing use of a Singer PIXL to pick up single clones. This work was supported by grants R01HG011676 to GS and SFL and R01AI164530 to SFL.

## Author Contributions

XL and SFL conceived the Bacterial Positioning System. XL developed the bacterial conjugation and recombination system. XL and WL developed the positional barcode system and performed the proof-of-concept experiments. WL performed the oligonucleotide and ORF demultiplexing experiments. WL developed protocols for ONT long read sequencing. DM developed the BPS analysis pipeline. LT, LC, and BP generated the *E. coli* ORF library. WL, DM, XL, SFL analyzed the data and wrote the paper. All authors edited the paper. GS and SFL secured grant funding.

## Supplemental Materials

**Table S1. Cost estimates on sequence verification, high throughput \***

**Table S2. Cost estimates on sequence verification, high throughput \***

**Table S3.**
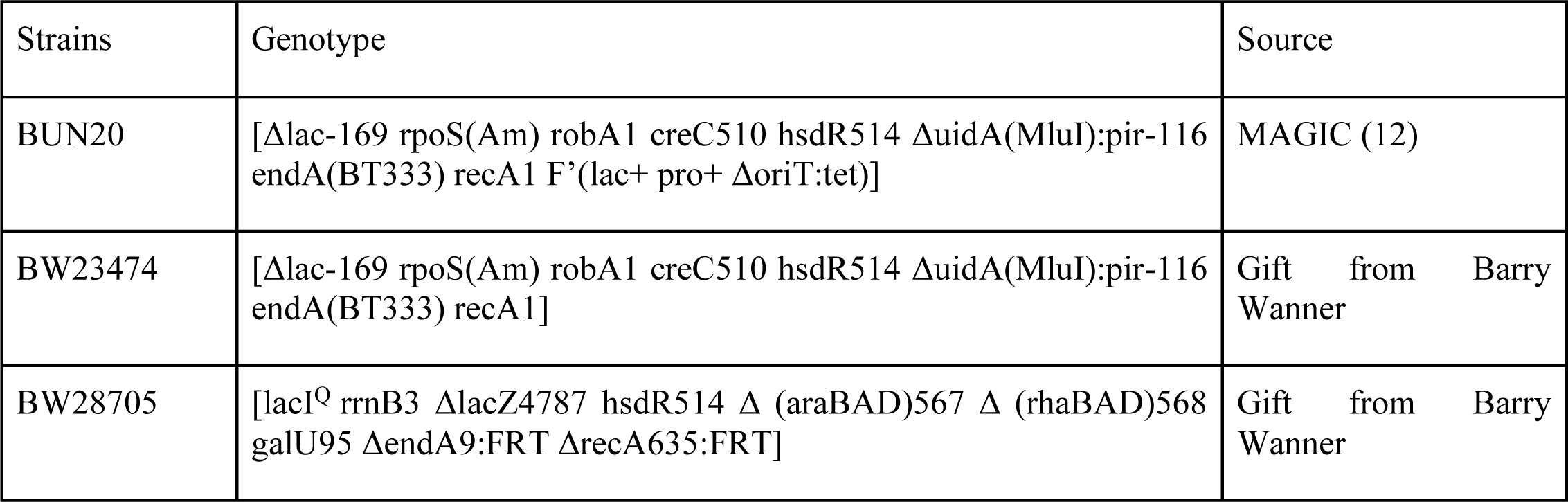
Bacterial strains used in this study.

**Table S4:** Plasmids used in *in vivo* barcoding.

| Name | Plasmid Type | Homology Regions | Swapping cassette | Other features | Source |
| --- | --- | --- | --- | --- | --- |
| pSL361 | Helper | NA | NA | P<sub>rhaBAD</sub>-I-SceI, P<sub>lac</sub>-red, recA | this paper |
| pSL937 | Recipient | HU, HD | P<sub>rhaBAD</sub>-relE | GmR | this paper |
| pSL438 | Donor | HU, HD | HygR-SacB | oriT, KanR | this paper |
| pSL438_BC | Donor | HU, HD | HygR-SacB | oriT, KanR, 15-nt barcodes | this paper |
| pSL439 | Donor | HU, HD | HygR-SacB | oriT, KanR | this paper |
| pSL439_BC | Donor | HU, HD | HygR-SacB | oriT, KanR, 15-nt barcodes | this paper |
| pSL1071 | Donor | HU, HD | NsrR-PheS | oriT, KanR | this paper |
| pSL1064 | Donor | HU, HD | HygR-SacB | oriT, KanR | this paper |
| pML104 | Helper | NA | NA | P<sub>lac</sub>-red, recA, SpeR | MAGIC (12) |

**Table S5.**
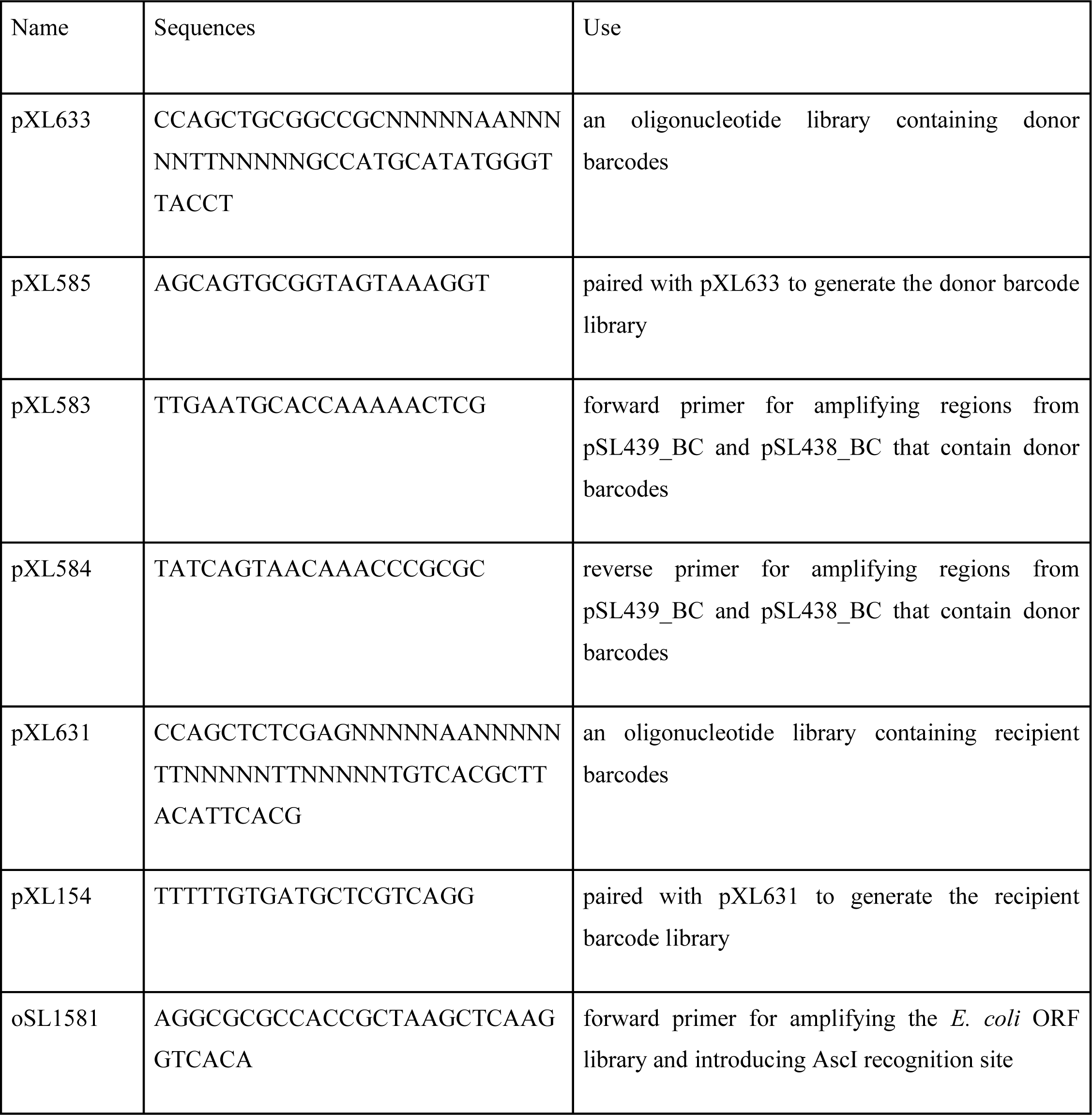
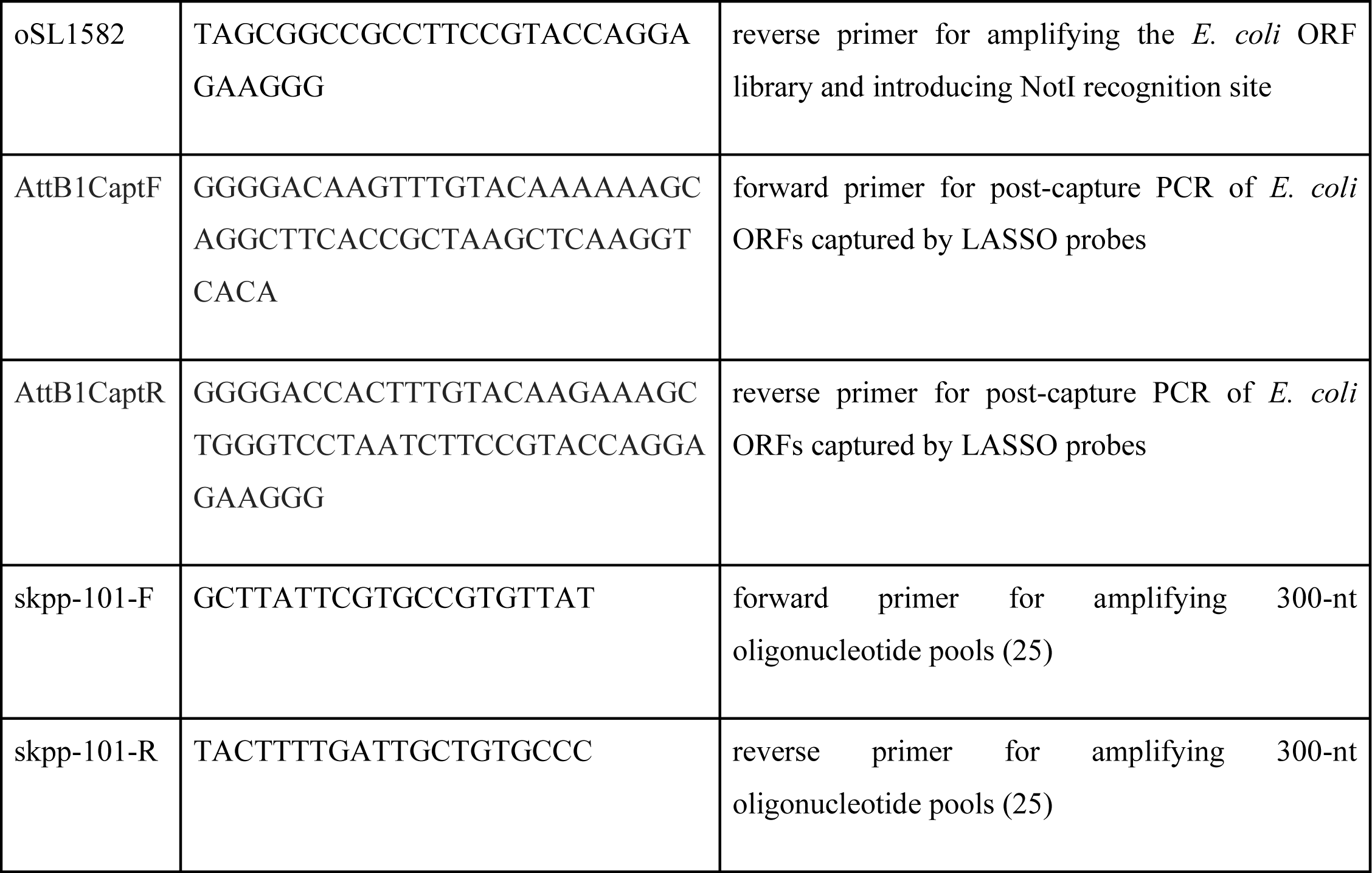
DNA oligonucleotides used in this study.

**Table S6. Sequences for 1,100 oligonucleotides randomly sampled from the human reference genome \***

**Table S7.** The composition of gap filling mix used for capturing *E. coli* ORFs.

| Component | Volume for 100 µL |
| --- | --- |
| PCR grade water | 73.8 µL |
| Glycerol | 10 µL |
| 10 mM dNTPs | 0.4 µL |
| Ampligase DNA Ligase (100 U µl<sup>-1</sup>) | 1 µL |
| Kapa HiFi | 0.8 µL |
| β-Nicotinamide adenine dinucleotide (NAD<sup>+</sup>) 50mM | 4 µL |
| 10× Ampligase DNA Ligase Buffer | 10 µL |

**Table S8. LASSO probes used in this study \***

**Table S9. Oligonucleotides used for Illumina sequencing \***

(* Tables S1, S2, S6, S8 and S9 are included in another uploaded file.)

**Figure S1. The recovery rate for donor DNA blocks (barcodes) at each position on 96-position arrays.**

**Figure S2. ONT read coverage across plates.**

**Figure S3. Sanger sequencing traces of 24 donor clones with purity scores ranging between 0.4 and 1.0.**

**Figure S4. The profile of base substitutions detected in the IDT and Twist oligonucleotide pools.**

**Figure S5. Abundance changes during cloning for the same subset of 100 oligonucleotide designs when large pools are parsed.**

