## Supplementary Figures for "Arrayed *in vivo* barcoding for multiplexed sequence verification of plasmid DNA and demultiplexing of pooled libraries"

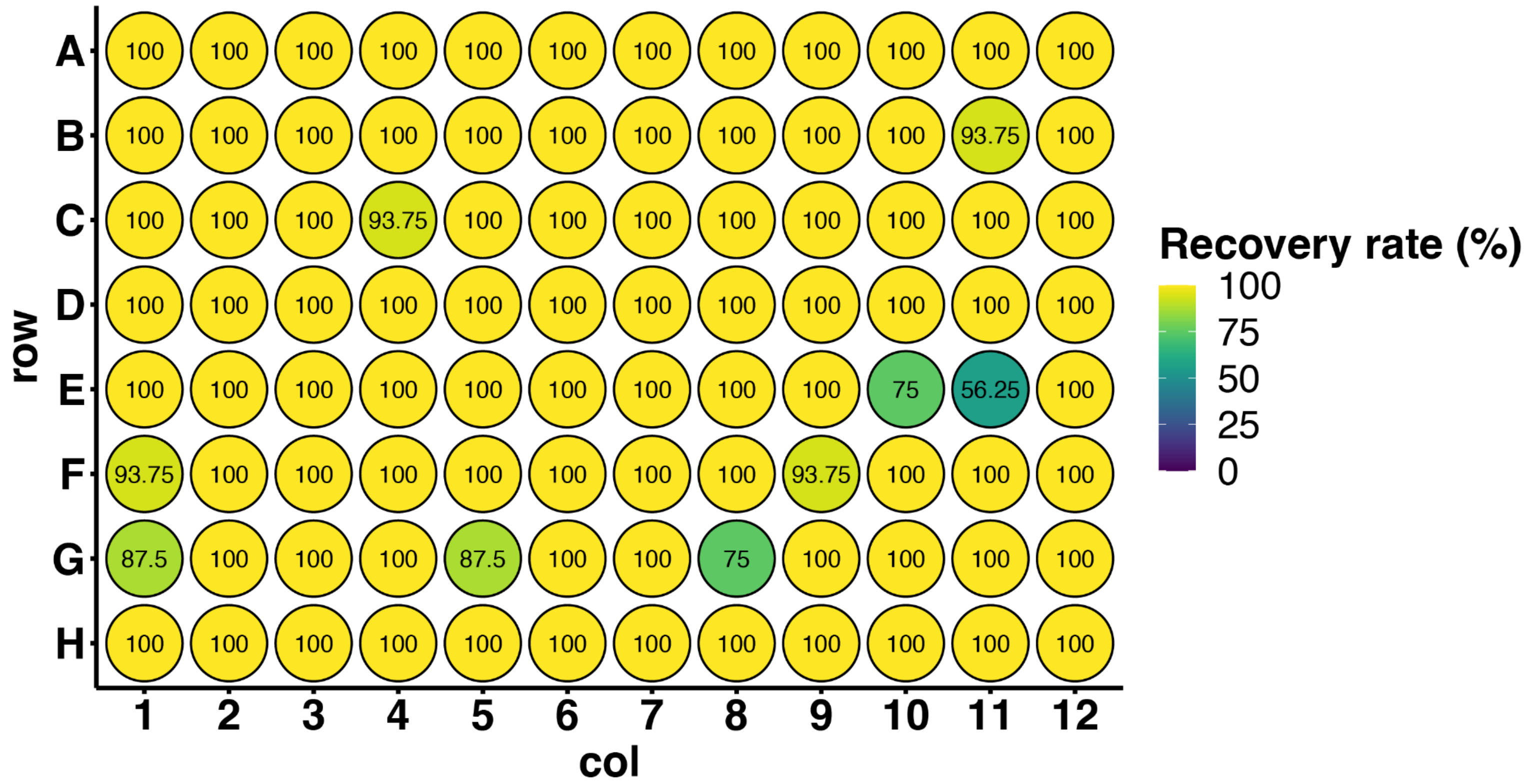


**Figure S1. The recovery rate for donor DNA blocks (barcodes) at each position on 96-position arrays.** 2× 96-arrayed donor barcode plates were each mated with 8× 96-arrayed recipient barcode plates. The per position recovery rate indicates the percentage of mating events for which one donor barcode can be accurately identified. The failures were, in all cases, due to a lack of sufficient sequencing coverage instead of inaccurate identification.

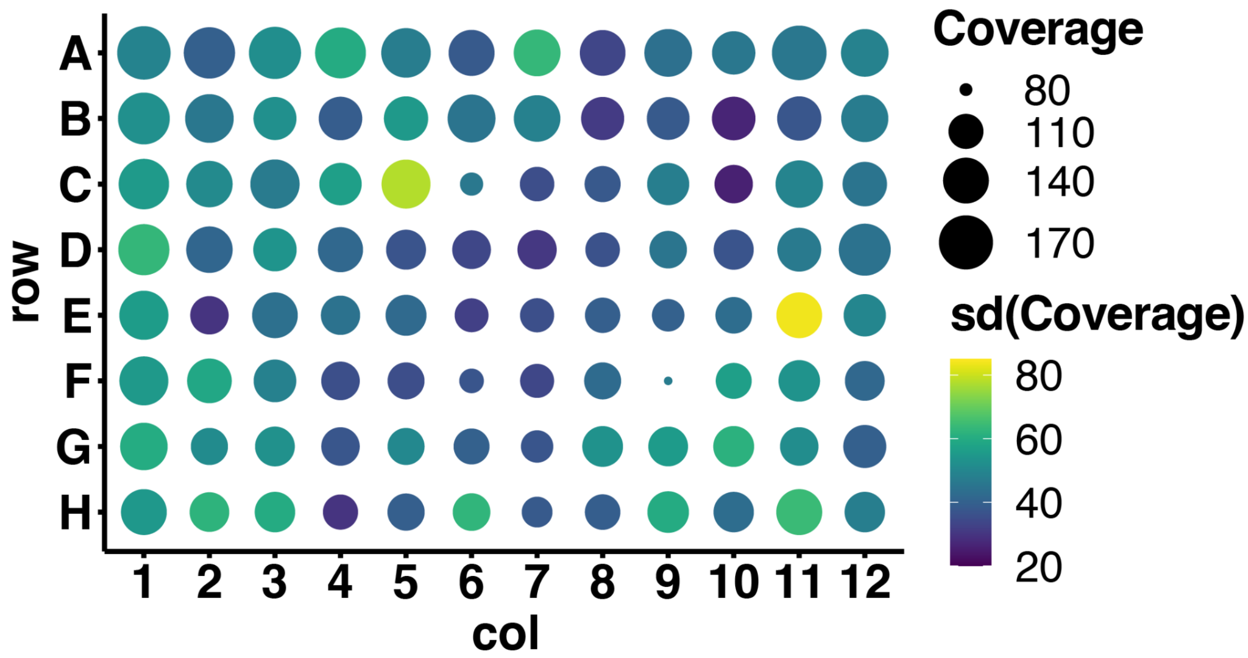


**Figure S2. ONT read coverage across plates.** The mean (size) and standard deviation (color) of ONT read coverage at each position from all plate matings. Some mating events result in insufficient number of ONT reads to detect the donor barcode and these lower the recovery rate. F9 is low in reads coverage (79.2) presumably because of a low recombination rate of the donor barcode clone (pSL438_BC) at this position. Peripheral positions on the array tend to have higher coverage because of the tendency of peripheral recombinant clones to generate large colonies, resulting in higher yields during a miniprep.

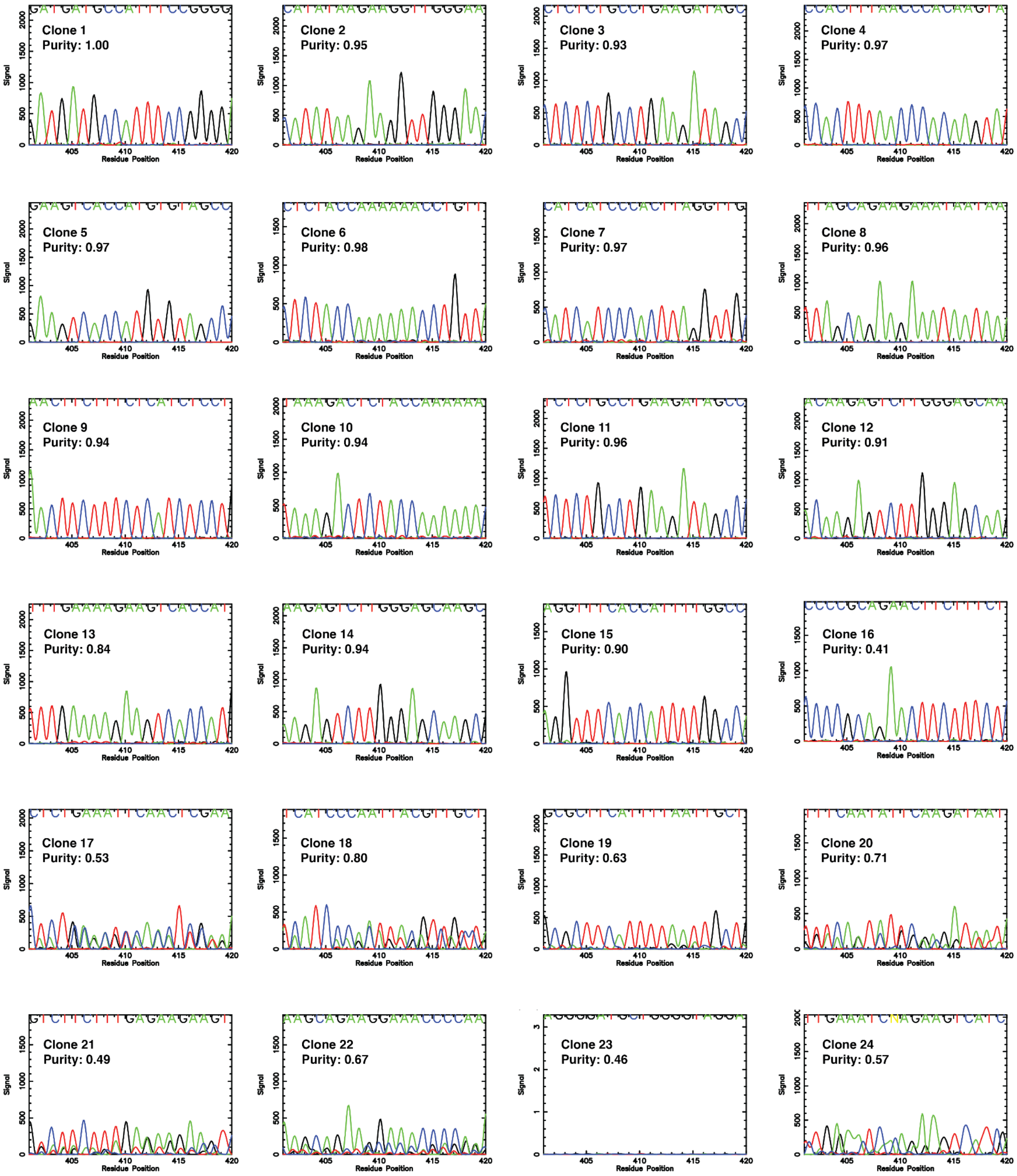


**Figure S3. Sanger sequencing traces of 24 donor clones with purity scores ranging between 0.4 and 1.0.** Colony PCR was performed with a pair of primers that anneals to the backbone of the donor plasmid. Amplicons were column purified and Sanger sequenced. Traces displayed here correspond to a 20-nt window of the DNA block (244 nt oligonucleotide design) being Sanger sequenced. Clones 1-15, with purity scores >0.8, have clean traces. Clone 16 has a purity score <0.8 but the trace suggests this clone is pure. Clones 17-24, with purity scores <0.8, show mixed traces. Clone 23 failed in Sanger sequencing.

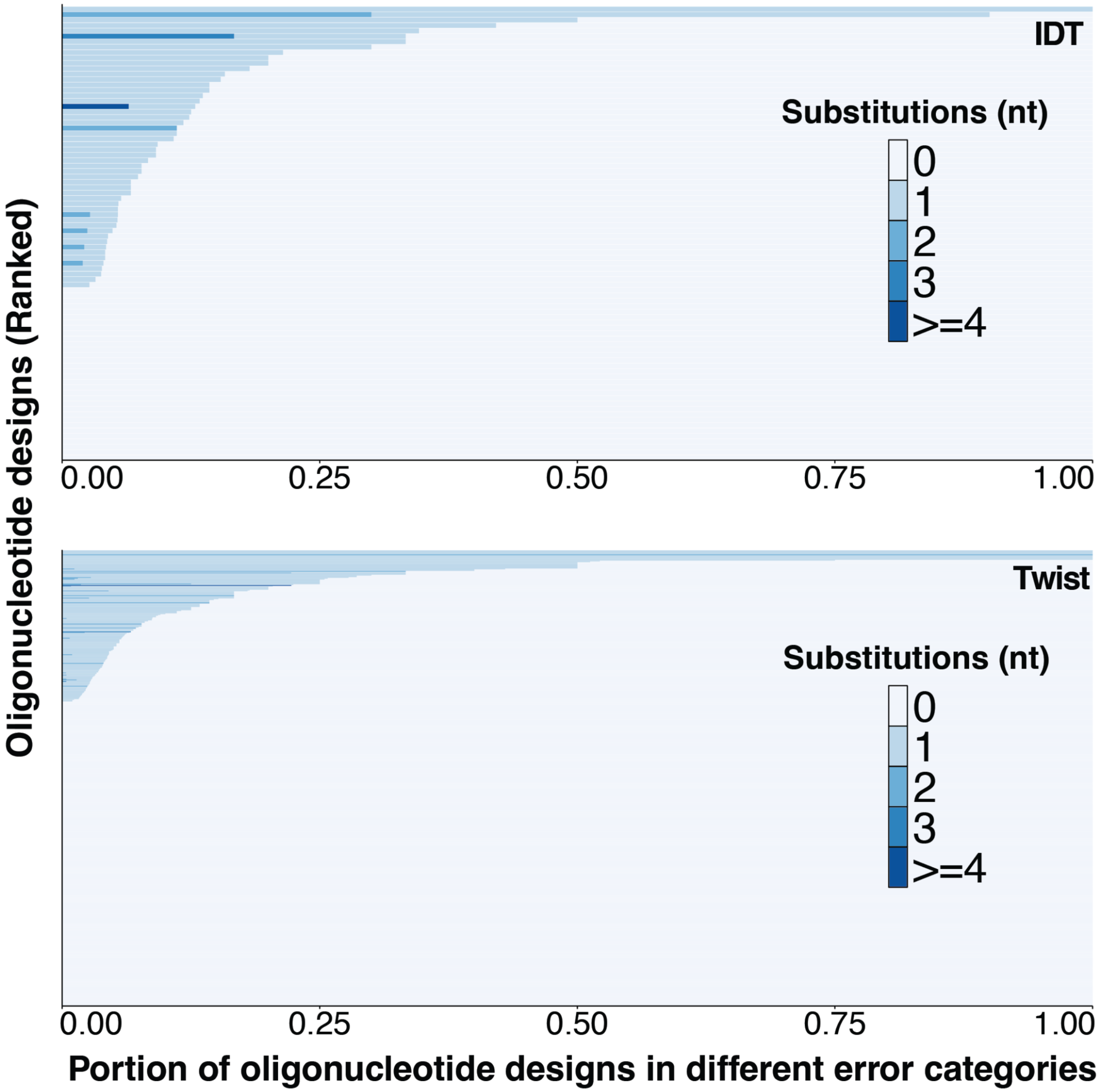


**Figure S4. The profile of base substitutions detected in the IDT and Twist oligonucleotide pools.** We focused on the full-length oligonucleotide designs that can be amplified and have been sequence-validated by BPS: a total of 85 and 580 oligonucleotide designs from IDT and Twist pools, respectively. The estimated base substitution rates are $4.75 \times 10^{-4}$/nt and $3.81 \times 10^{-4}$/nt for IDT and Twist pools, respectively.

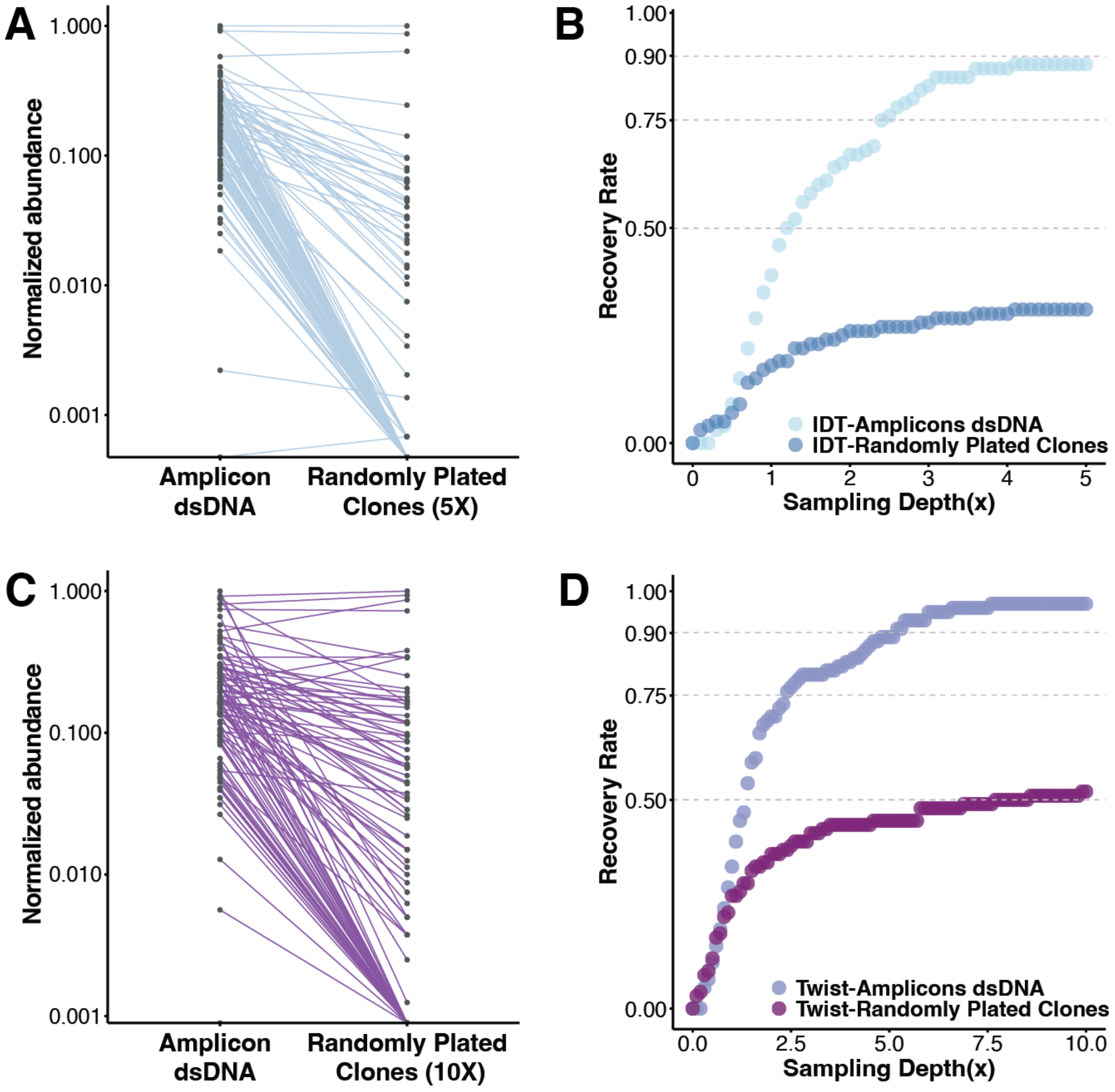


**Figure S5. Abundance changes during cloning for the same subset of 100 oligonucleotide designs when large pools are parsed.** To determine whether an increased size and complexity of a DNA library may impact the recovery rate, we ordered all of the 1,100 oligonucleotide designs from both IDT and Twist and cloned them as pools into the BPS donor plasmids and cells following the same amplification, ligation, and transformation protocols as described in Figure 2. We show results for the small IDT pool containing the same 100 oligonucleotide designs in Figure 2D. (A) and (C) The change in normalized abundance in (A) the IDT and (C) the Twist pools. (B) and (D) The recovery rate at different sampling depths for (B) the IDT and (D) the Twist pools. The full data set for the Twist pool, including all 1,100 oligonucleotide designs, can be found in Figures 2F and 2G.
